# Organoid Models Established from Primary Tumors and Patient-Derived Xenograft Tumors Reflect Platinum Sensitivity of Ovarian Cancer Patients

**DOI:** 10.1101/2024.06.28.601283

**Authors:** Parisa Nikeghbal, Dorsa Zamanian, Danielle Burke, Mara P. Steinkamp

## Abstract

**Background:** Ovarian cancer (OC) remains the deadliest gynecological cancer, primarily due to late-stage diagnosis and high rates of chemotherapy resistance and recurrence. Lack of representative preclinical models complicate the challenges of discovering effective therapies, especially for platinum-resistant OC. Patient-derived xenograft (PDX) models maintain the genetic characteristics of the original tumor and are ideal for testing candidate therapies *in vivo*, but their high cost limits their feasibility for high-throughput drug screening. Organoid models mimic the tumor’s 3D structure and preserve intra-tumoral heterogeneity. While organoids established directly from primary patient tumors are the optimal model for personalized drug response studies, the supply of primary tissue is often limited. Patient-derived xenograft tumors can be passaged in mice and provide a renewable source of cancer cells for organoids. This study aimed to determine if PDX-derived organoids (PDXOs) can reflect patient responses to chemotherapy similarly to primary patient-derived organoids (PDOs).

**Methods:** 3D tumor organoid cultures were established from paired primary OC and PDX samples. Organoid viability after 72-hour treatment with paclitaxel (PTX), carboplatin (CBDCA), or their combination was compared between organoids derived directly from the patient or from the PDX models. The *in vitro* drug responses of PDXOs and PDOs were then compared to defined patient clinical responses: platinum-sensitive (initial response to standard platinum/taxol therapy lasting >6 months post-treatment), platinum-resistant (initial response to standard chemotherapy lasting < 6 months), or platinum-refractory (no initial response to standard chemotherapy).

**Results:** In drug response assays, PDXOs and PDOs demonstrated similar sensitivity to standard chemotherapy and also reliably reflected patient responses based on the clinical designation of platinum sensitivity. While organoids derived from the ascites were smaller with a denser morphology, their drug response mirrored that of the organoids derived from solid tumor. Platinum-sensitive cases exhibited significant reductions (around 50% reduction) in organoid viability when treated with carboplatin, paclitaxel, or their combination. Platinum-resistant or refractory organoids showed little to no reduction in viability with carboplatin or paclitaxel monotherapy or the combination. Organoids derived from one platinum-resistant case did show a small but significant reduction in viability with single-agent paclitaxel, suggesting that organoid models might predict response to second-line paclitaxel therapy.

**Conclusion:** This study demonstrates that PDXOs respond to drugs similarly to PDOs and confirms that both models effectively mirror patient response to standard chemotherapy. This highlights the potential of PDXOs as renewable models for screening novel therapies and developing personalized strategies in OC.

**SIMPLE SUMMARY:** Ovarian cancer (OC) remains the most lethal gynecological cancer, largely due to its late diagnosis and resistance to chemotherapy. In order to identify novel therapies to treat ovarian cancer, we need better *in vitro* models that represent the genetic heterogeneity of the patient population. This study evaluates patient-derived organoid models established either directly from patient samples (PDOs) or from patient samples that were first passaged in mice and are referred to as patient-derived xenograft organoids (PDXOs). For each patient, organoid response to standard chemotherapy based on organoid viability assays was compared to the patient’s clinical designation of platinum sensitivity, which is categorized as platinum-sensitive, platinum-resistant, or platinum-refractory based on their response to standard chemotherapy. We demonstrated that both PDOs and PDXOs accurately reflect the patient’s clinical designation, suggesting their utility as effective models for testing new therapies and personalizing treatment. Importantly, by providing a renewable source of patient-derived cells, PDXOs extend the utility of each sample, making organoids essential tools for developing and refining personalized treatment strategies in oncology.

## 1. INTRODUCTION

Ovarian cancer (OC) has one of the highest mortality rates among gynecological cancers, largely due to its late-stage diagnosis, resistance to chemotherapy, and recurrence after first line treatment [1]. OC is typically treated with surgical debulking and platinum-based chemotherapy [2] with a 70-80% initial response rate and a 50% complete remission [3]. Despite these efforts, over 80% of patients recur with resistant disease, underscoring the need to identify effective second-line treatment options [2, 4]. The platinum-free interval (PFI) classifies patients into platinum-sensitive (PFI ≥ 6 months), platinum-resistant (PFI < 6 months), and platinum-refractory groups, each with varying outcomes: those in the sensitive group have a median overall survival (OS) of 24–36 months, whereas resistant and refractory groups face worse prognoses, with OS of 9–12 months and 3–5 months, respectively [5-8]. OC is a highly heterogeneous disease with a critical shortage of preclinical models that accurately reflect patient variability, underscoring the urgent need for better models to identify the most effective treatment for each patient [9]. Patient-derived xenograft (PDX) models maintain the genetic diversity and morphological characteristics of the original tumors and can be used to test drug response *in vivo* [10-13]. However, their use is hindered by high costs and lengthy timelines, which limit their utility in large-scale drug screening [14-16].

Traditionally, the majority of OC drug testing has relied on the response of OC cell lines grown as monolayers. These cell lines are known to lose their heterogeneity due to rapid clonal selection [17-21]. Organoid cultures that replicate the three-dimensional complexity of the tumor microenvironment and the genetic diversity of OC tumors can be established from digested tumor tissue [22, 23]. Organoids closely recapitulate the tumor morphology and preserve intra-tumoral heterogeneity [16, 24], unlike two-dimensional cultures of immortalized cancer cell lines. Organoids not only closely mimic clinical drug responses but also have the potential to promote personalized therapy by reflecting patient-specific responses and revealing the genetic underpinnings of drug resistance [25].

Studies directly comparing responses in colorectal and metastatic gastrointestinal cancer organoids to patient response have shown that these *in vitro* models can effectively reflect patient responses to both chemotherapy and targeted therapies [26, 27]. High-throughput studies further validate the organoids’ utility in mapping drug efficacy, showing strong correlations with clinical outcomes in OC [8, 16, 25, 28, 29].

Organoid drug response assays have even been used to test advanced drug response predictions via neural networks [30]. Moreover, these organoid assays streamline the drug testing process, aligning it with therapeutic decision-making by rapidly identifying patient-specific drug sensitivities [31, 32]. One previous study anticipated carboplatin (CBDCA) resistance in OC by combining organoid drug response data with gene expression analysis to forecast outcomes [8].Our study explores the utility of PDX-derived organoids (PDXOs) for high throughput drug screening in organoid models. This approach will help identify potentially effective therapies, which we can then test in paired PDX models, creating an *in vitro* to *in vivo* pipeline. Intraperitoneal (IP) engraftment of cancer spheroids isolated from OC primary malignant ascites and engrafted into NSG mice models OC peritoneal dissemination. These PDX can then be used to generate organoids from both solid tumor and ascites samples. PDXOs derived from breast cancer models have accurately predicted therapeutic responses *in vivo* [33-35]. A previous study in ovarian clear cells carcinoma demonstrated that patient-derived tumor organoids and xenografts could replicate the clinical drug resistance observed in patients [36]. However, the correlation in response between patients, PDOs, and PDXOs has not been directly compared. We have established ovarian cancer PDX models from the malignant ascites of a cohort of high-grade serous OC cases. Clinical response data for first-line carboplatin/paclitaxel chemotherapy was collected for these patients. Here we have compared the drug responses among PDOs derived from patient ascites and PDXOs derived from both PDX ascites and solid tumors, to corresponding patient clinical outcomes. Our results indicate that PDXOs derived from ascites or solid tumor, similar to primary PDOs, consistently reflect the clinical response to the standard first-line chemotherapy. This correlation underscores the potential of 3D *in vitro* OC organoids derived from PDXs as an important tool for initial high throughput screening. PDX organoids support the growth of patient-derived cancer cells in a manner that closely mimics the *in vivo* environment and allow us to directly translate drug response from *in vitro* to *in vivo* models with the goal of representing the clinical response.

## 2. MATERIAL AND METHODS

### Isolation of cancer cells from patient samples

Malignant ascites fluid was collected from consenting OC patients undergoing cytoreductive surgery at the University of New Mexico Comprehensive Cancer Center as detailed in Steinkamp et al, 2023 [13]. Acquisition of patient samples was approved by the UNM Health Science Center Institutional Review Board (protocol no. INST1509) and studies were conducted in accordance with the U.S. Common Rule. Red blood cells were lysed using Ammonium Chloride Solution (STEMCELL Technologies, 07800). Cancer spheroids were isolated from the ascites fluid, cryopreserved in freezing media (5% DMSO/95% FBS, Gibco, Thermo Fisher Scientific), and stored at -80°C prior to processing for organoid development.

### Patient-derived xenografts

Fresh or cryopreserved spheroids isolated from the ascites fluid of OC patients were injected into the peritoneal cavity of in-house bred NSG mice (RRID: IMSR_JAX:005557) to establish orthotopic PDX models of disseminated OC, as detailed in Steinkamp et al, 2023 [13]. PDX-engrafted mice were euthanized at a humane endpoint when ascites fluid accumulation caused abdominal distention or when mice showed signs of wasting. At necropsy, solid tumors and ascites fluid were collected and cryopreserved in 5% DMSO/95% FBS. PDXOs were generated from cryopreserved PDX solid tumors and ascites.

### Organoid Formation

Development of patient-derived cancer organoids (PDOs) or PDX cancer organoids (PDXOs) followed protocols adapted from [37, 38]. Briefly, solid tumors were mechanically dissociated then incubated for 30 minutes at 37°C with rotation in digestion buffer (human wash buffer (DMEM/F12 (Gibco, 21041025), 10 mM HEPES (Gibco, 15-630-080), 1X GlutaMAX (Gibco, 35-050-061), 100 µg/ml primocin (Invivogen, ant-pm-1), and 0.1% BSA (Sigma, A8531) supplemented with 2 U/mL dispase II (Sigma, D4693), 1 mg/mL collagenase P (Millipore, 11213857001), and 2 µL/ml DNase I (New England Biolabs, M0303S)). Digested tissue was strained through a 40 µm strainer to obtain a single-cell suspension, washed twice with human wash buffer, and centrifuged at 500 x g for 10 minutes after each wash. Primary ascites-derived cancer spheroids were centrifuged at 500 x g and resuspended in digestion buffer.

To establish cancer organoid cultures, 50,000 cancer cells/well were embedded in 100% ultimatrix (Cultrex, BME001-10) in 50 µL domes in a 24-well plate. Domes were overlaid with 450 µL of organoid media (Advanced DMEM/F12 (Gibco, 12634028) supplemented with 1X GlutaMAX, 10 mM HEPES, and Pen/Strep, 100 ng/ml recombinant human noggin (Peprotech, 120-10C), 250 ng/ml R-spondin-1 (R&D Systems, 4645-RS-025), 100 ng/ml FGF10 (Peprotech, 100-26), 37.5 ng/ml Heregulin beta-1 (US Biological, H2030-50), 50 ng/ml EGF (R&D Systems, 236-EG), B27 supplement (1:50, Thermofisher, 17504044), 1.25 mM N-acetylcysteine (Sigma, A9165), 100 nM ß-estradiol (Sigma, E2758), 2% primocin, 5 mM nicotinamide (Sigma, N0636), 5 µM A83-01 (Tocris, 2939), 10 µM Y-27632 (Sigma, Y0503),10 µM forskolin (Tocris, 1099), and 250 µg/ml hydrocortisone (Sigma, H0888). Formation of organoids was monitored on an Incucyte S3 Live Cell Analysis instrument with images acquired every 12 hours. Phase contrast images are shown in supplementary figures. Initial organoid formation from single cells took 6-14 days.

### Drug Screening

Drug screening was performed in 96-well white plates (CELLSTAR, Sigma, M1062-40EA) with µClear® bottoms. Each well was seeded with 20-30 organoids in ultimatrix/organoid media in a 1:1 ratio and seeded as 5 µL domes following the Thiel lab’s protocol [38]. After dome solidification, 200 µL of pre-warmed organoid culture media was added to each well. The following day, the organoids were treated with standard chemotherapy (carboplatin, paclitaxel, or the combination) at concentrations specified in the figures. 0.1% DMSO diluted in organoid media served as the vehicle control. Treatment with 10 µM afatinib, a pan-ErbB receptor inhibitor, effectively kills ovarian cancer organoids and was used as a control for cell killing. After 72 hours, organoid cell viability was determined using the CellTiter-Glo® 3D Cell Viability Assay (Promega, G9682) following the manufacturer’s protocol. Briefly, an equal amount of CellTiter-Glo® 3D reagent was added to the media in each well and plates were agitated vigorously in a microplate reader for 5 minutes, then allowed to sit at room temperature for 25 minutes to stabilize the luminescence signal. Luminescence was measured using a BioTek Synergy Neo2 Multi-Mode high-sensitivity plate reader (**Figure 1**). All assays tested 6 or more wells/condition as technical replicates. Two independent drug response studies were run per PDO/PDXO as biological replicates.

**Figure 1.**
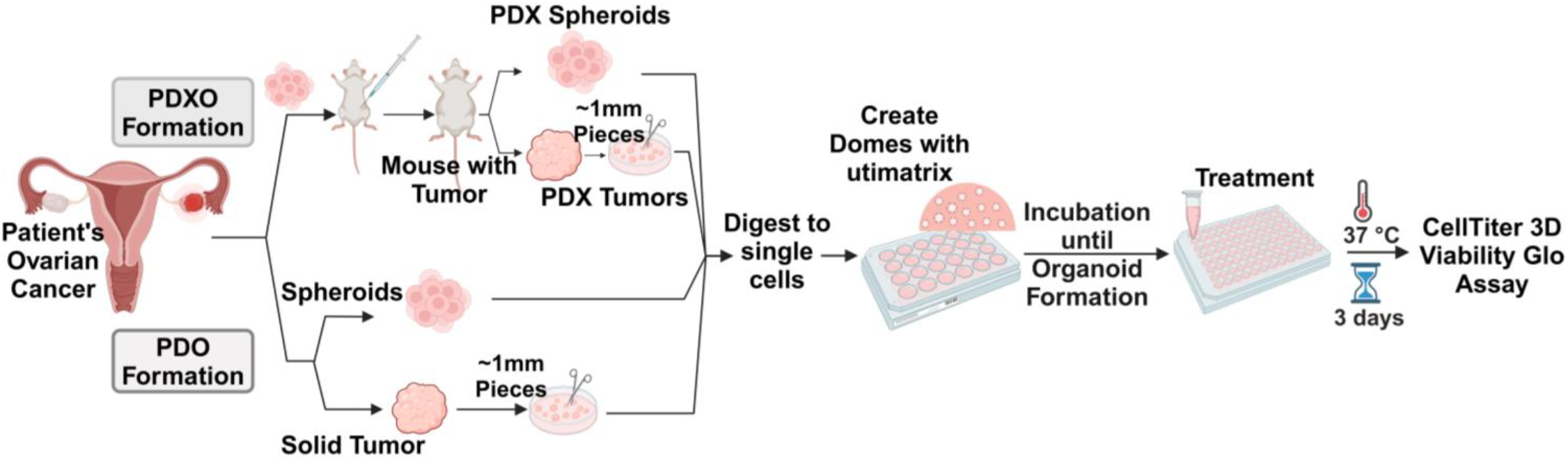
Comprehensive workflow for developing and evaluating chemotherapy responses in patient-derived and PDX-derived ovarian cancer organoids. A schematic of the generation and treatment of patient-derived organoids (PDOs) and patient-derived xenograft organoids (PDXOs) from ovarian cancer samples. Fresh primary ovarian cancer cells are first collected from the malignant ascites of patients. These cancer cells are engrafted into NSG mice to establish PDXs. Once tumors develop, the resulting ascites or solid tumors can be used for PDXO formation. For PDO formation, primary ascites or solid tumors are processed directly. To seed organoids, solid tumors are cut into ∼1 mm pieces, and all samples are enzymatically digested into single-cell suspensions. Cancer cells are subsequently embedded in ultimatrix, an organic extracellular matrix to form organoids. Once formed, organoids are harvested and seeded in 5 µL ultimatrix domes in 96-well plates. One day later, they are treated with chemotherapeutic agents CBDCA, PTX, or their combination. After three days of treatment, organoid viability is assessed using the CellTiter-Glo® 3D Cell Viability Assay. This figure was created with BioRender.com.

### Statistical Analyses

Statistical analysis was performed using GraphPad Prism. One-way ANOVA examined differences between groups. A p-value <0.05 was considered significant.

### Data Availability

The data generated in this study are available within the article and its Supplemental Data.

## 3. RESULTS

### Characterization of Chemotherapy Sensitivity and Growth Dynamics in Patient-Derived and PDX-Derived Ovarian Cancer Organoids

In order to develop patient-derived OC models to test between-patient variation in response to standard and targeted treatments, cancer cells isolated from primary OC malignant ascites, or from PDX solid tumors or malignant ascites, were seeded in matrix domes to form patient-derived organoids (PDOs) or PDX-derived organoids (PDXOs) respectively (**Figure 1**). Patient-derived organoid models have been shown to preserve the genetic and phenotypic characteristics of the original OC patient tumors [16, 30]. When seeded in a 96-well format, organoid cultures allow for rapid high-throughput testing of single agents and combination treatments *in vitro*.

For this study, PDOs and PDXOs were established from five high-grade OC patients (**Table 1**). The clinical designation of platinum sensitivity based on response to standard chemotherapy is listed. Samples are designated: platinum-sensitive (partial or complete response to standard carboplatin/paclitaxel treatment with recurrence >6 months after treatment), platinum-resistant (disease progression on therapy or recurrence within 6 months), or platinum-refractory (no response to initial treatment) [39-41].

**Table 1.** Clinical designation of ovarian cancer patients who donated the samples used in this study. Ovarian cancer patients are arranged by the clinical designation of their response to platinum-based chemotherapy. Patients are classified as platinum-sensitive (experiencing partial or complete response to standard CBDCA/PTX treatment with recurrence more than six months after treatment), platinum-resistant (showing disease progression on therapy or recurrence within six months), or platinum-refractory (no response to initial treatment).

| **Platinum sensitivity** | **Patient #** | **Origin of the Organoids** |
| --- | --- | --- |
| **Sensitive** | **Pt 18** | **Ascites PDO**<br>**Solid tumor & Ascites PDXO** |
|  | **Pt 20** | **Ascites PDO**<br>**Solid Tumor & Ascites PDXO** |
| **Resistant** | **Pt 2** | **Ascites PDO**<br>**Solid tumor & Ascites PDXO** |
|  | **Pt 80** | **Ascites PDO**<br>**Solid tumor & Ascites PDXO** |
| **Refractory** | **Pt 1** | **Ascites PDO**<br>**Solid Tumor & Ascites PDXO** |

Ascites-derived PDOs and PDXOs seeded as single cells suspensions formed small organoids within 7–14 days (**Supplemental Figure 1**) and 7–10 days (**Supplemental Figure 2**) respectively. In contrast, PDXOs derived from PDX solid tumors formed small organoids by 2–3 days and large organoids within 6–9 days (**Supplemental Figure 3**). Overall, both PDOs and PDXOs derived from ascites samples grew slower and formed smaller organoids compared to those derived from PDX solid tumors.

**Figure 2.**
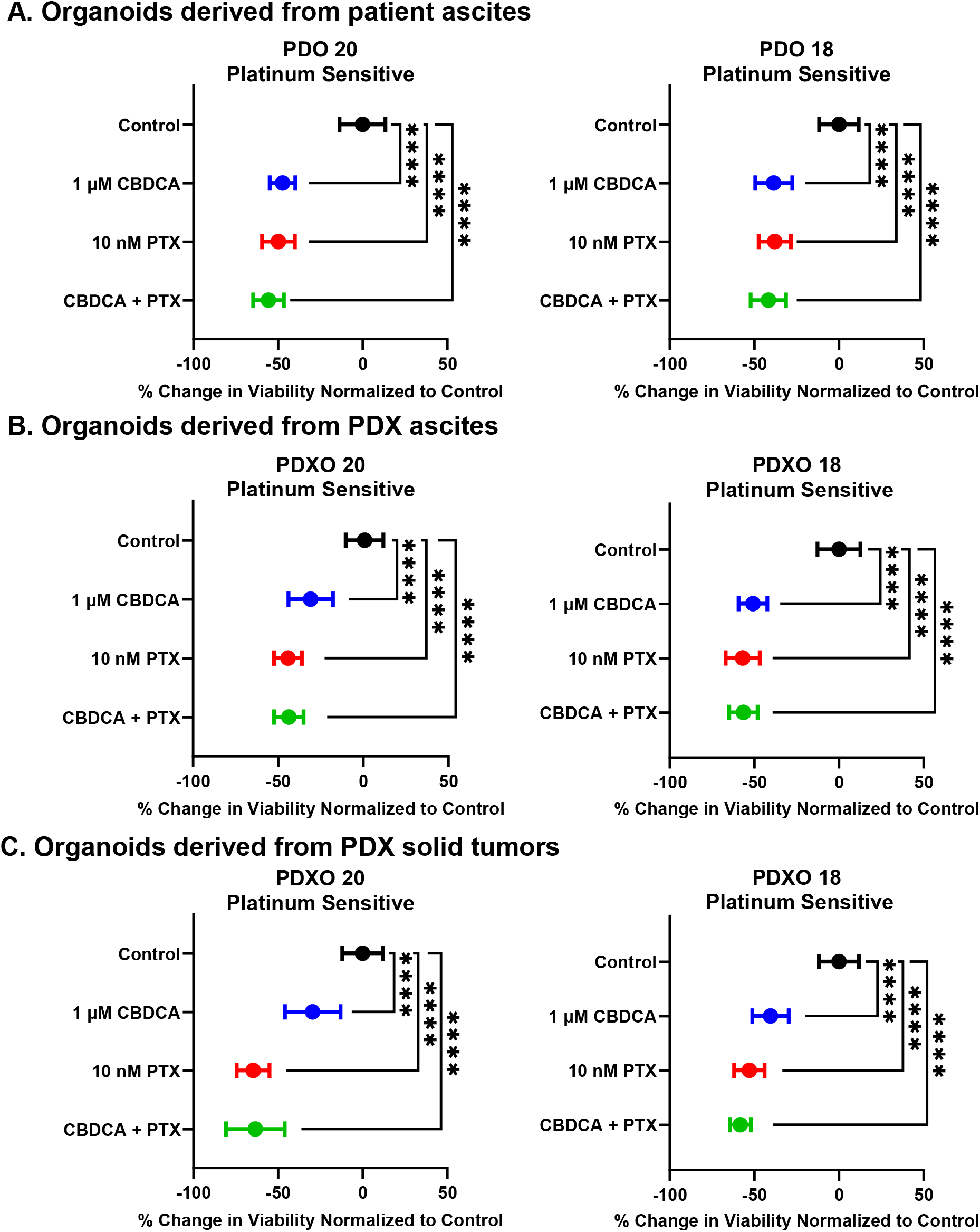
Organoids derived from platinum-sensitive patients are sensitive to CBDCA, PTX, and the combination. Each graph shows the change in cell viability relative to untreated control organoids (set to 0%) and organoids treated with CBDCA, PTX, and their combination after 72 hours of treatment. **A**. PDOs derived from the malignant ascites of platinum-sensitive patients. **B**. PDXOs derived from PDX ascites of the same patients as A. **C**. PDXOs derived from PDX solid tumors from the same patients as in A. The graphs display the difference in organoid viability, assessed by the CellTiter 3D Glo Assay, normalized to untreated control. Negative values indicate a decrease in viability, which suggests response to treatment. Error bars represent standard deviation, based on two biological replicates with 6 or more technical replicates per experiment.. Statistical significance is marked by asterisks (ns: not significant; * p<0.05; ** p<0.01; *** p<0.001; **** p<0.0001), assessed via one-way ANOVA.

**Figure 3.**
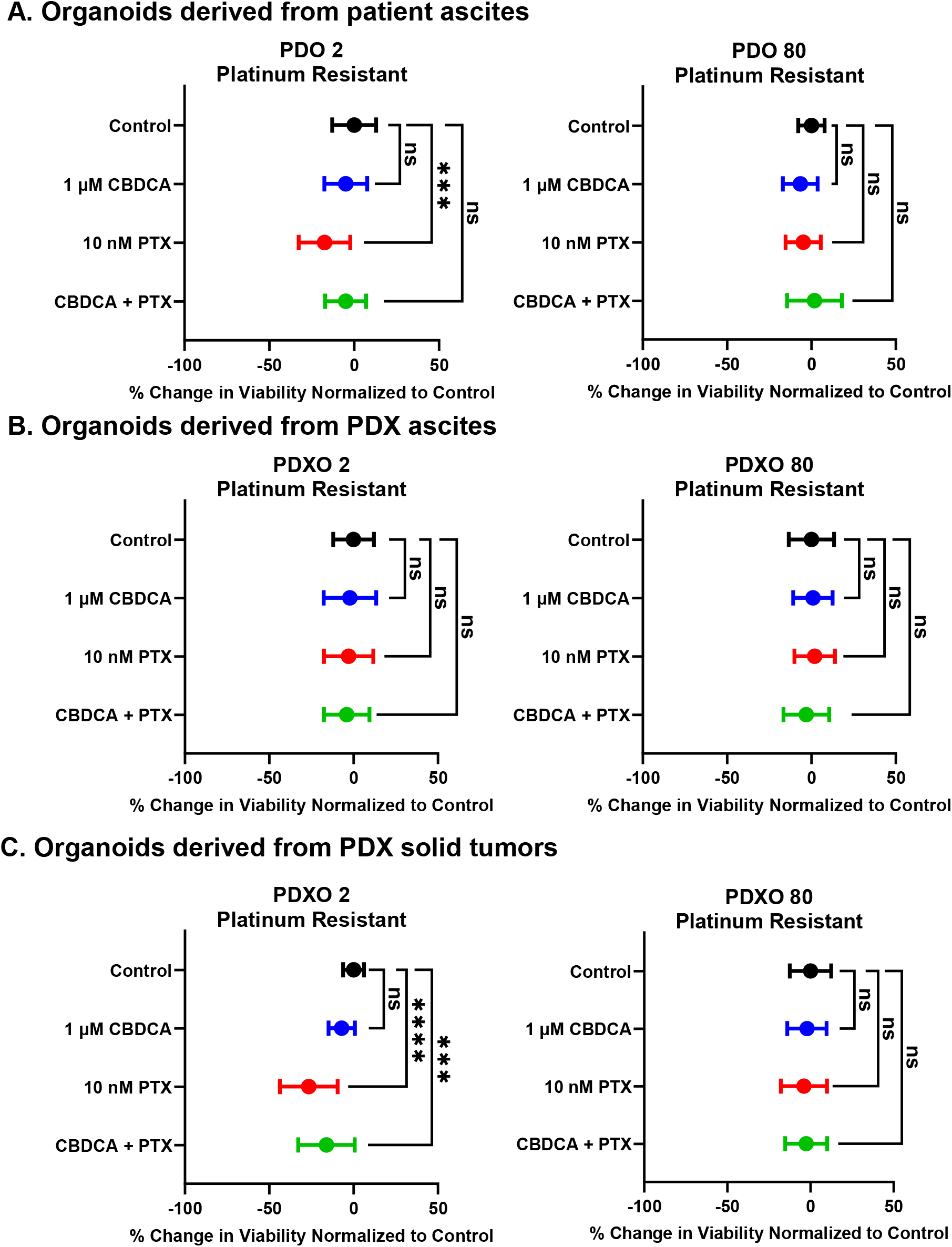
Platinum-Resistant Models Demonstrate Resistance to Combination Therapy. Each graph displays the change in cell viability relative to untreated control organoids (set to 0%) and organoids treated with CBDCA, PTX, and their combination after 72 hours of treatment. **A**. PDOs derived from malignant ascites of platinum-resistant patients. **B**. Paired PDXOs derived from ascites. **C**. Paired PDXOs derived from solid tumors. The graphs display the difference in organoid viability, assessed by the CellTiter 3D Glo Assay, normalized to untreated control. More negative values indicate response to treatment. Error bars represent standard deviation based on two biological replicates with 6 or more technical replicates per experiment. Statistical significance is marked by asterisks (ns: not significant; * p<0.05; ** p<0.01; *** p<0.001; **** p<0.0001), assessed via one-way ANOVA.

To assess organoid response to standard first line chemotherapy, PDOs and PDXOs were plated in 5 µl domes in 96 well plates and treated with 1 µM carboplatin, 10 nM paclitaxel, or the carboplatin/paclitaxel combination for 72 hours. Cell viability was assessed using a 3D luminescent viability assay that quantifies ATP levels. As a control for effective cancer cell killing, organoids were treated with 10 µM afatinib, a pan-ErbB receptor inhibitor that effectively kills ovarian cancer organoids regardless of their platinum sensitivity. PDO models yielded lower luminescence than PDXOs. However, luminescence intensity was consistent among PDOs derived from the five patients. Across all models, untreated organoids demonstrated luminescence within the linear range of the assay and showed a significantly reduced luminescence with afatinib treatment (**Supplemental Figure 4**).

**Figure 4.**
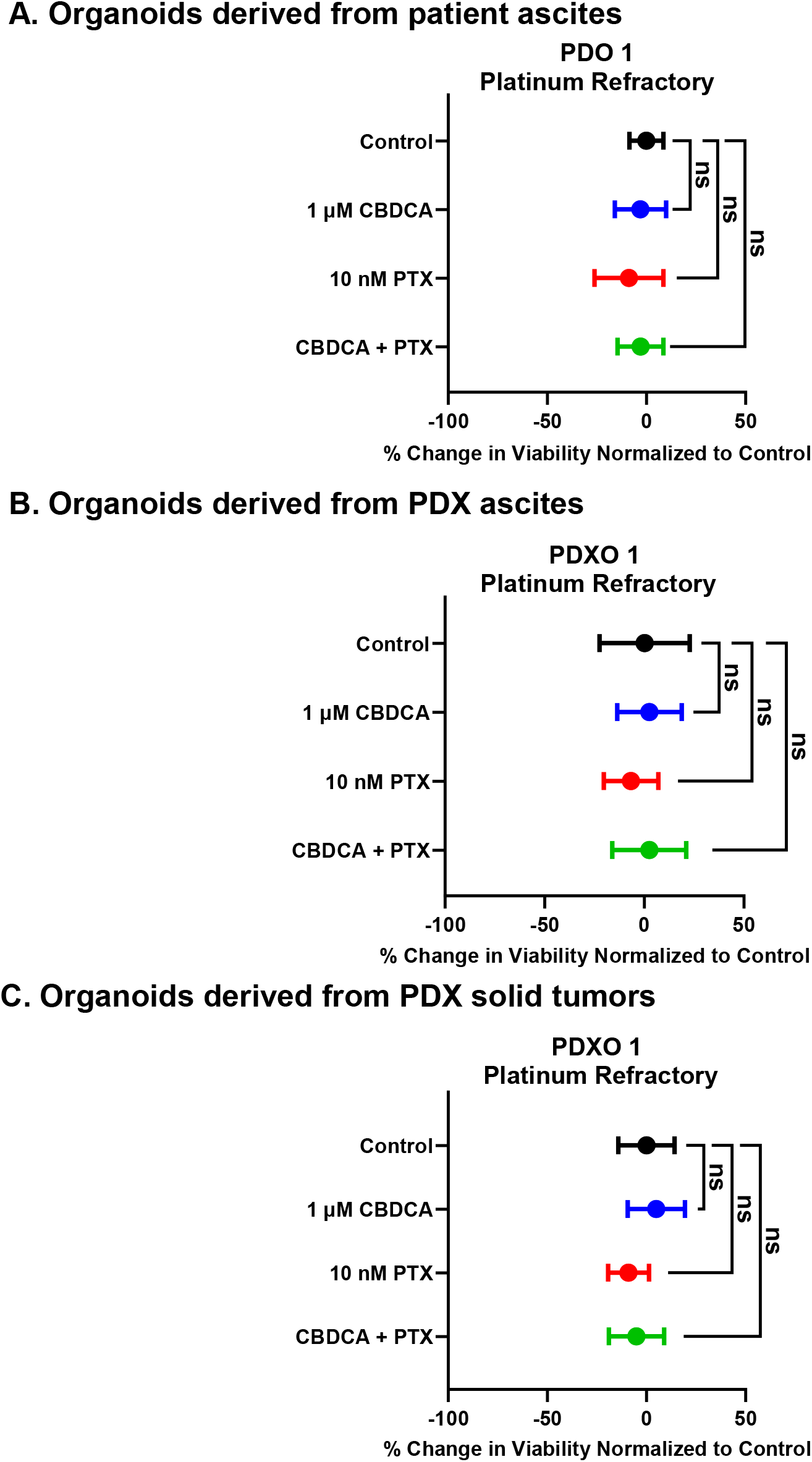
Platinum-Refractory Models Show Resistance to Treatment. Each graph displays the change in cell viability relative to untreated control organoids (set to 0%) and organoids treated with CBDCA, PTX, and their combination after 72 hours of treatment. **A**. PDOs derived from malignant ascites of platinum-refractory patients. **B**. Paired PDXOs derived from ascites. **C**. Paired PDXOs derived from solid tumors. The graphs display the difference in organoid viability, assessed by the CellTiter 3D Glo Assay, normalized to untreated control. More negative values indicate response to treatment. Error bars represent standard deviation based on two biological replicates with 6 or more technical replicates per experiment. Statistical significance is marked by asterisks (ns: not significant; * p<0.05; ** p<0.01; *** p<0.001; **** p<0.0001), assessed via one-way ANOVA.

### Platinum-Sensitive Models Show High Predictability in Chemotherapy Response

For comparison of treatment response across organoid models and across experiments, the change in organoid viability with treatment was normalized to average viability of the untreated controls (0% change). PDOs 20 and 18, derived from malignant ascites of platinum-sensitive patients, exhibited significant reductions in viability when treated with 1 µM carboplatin, 10 nM paclitaxel, or their combination. In both cases, combination therapy led to the most pronounced decrease in cell viability, though single-agent treatments also demonstrated substantial effectiveness. These results are consistent with the clinical platinum sensitivity of these patients (**Figure 2A**). Similarly, PDXO models from the same patients (PDXO 20 and PDXO 18), whether derived from ascites (**Figure 2B**) or from solid tumors (**Figure 2C**), mirrored the responses of their PDO counterparts, showing comparable reductions in viability across all treatment conditions. The close alignment of PDO and PDXO responses—whether from ascites or solid tumors—confirms that these models retain the characteristics of the patient tumor in platinum-sensitive cases regardless of the difference in morphology between the PDOs and PDXOs.

### Platinum-Resistant Models Demonstrate Resistance to Combination Therapy

PDOs derived from platinum-resistant cancers exhibited resistance to carboplatin alone and to the combination carboplatin/paclitaxel as expected based on their clinical response. Combination treatment in PDO 80 even led to a slight but significant increase in organoid viability above that of untreated control (**Figure 3A**). However, response to paclitaxel varied between the two models. PDO 2, but not PDO 80, displayed a small but significant reduction in cell viability when treated with paclitaxel alone (**Figure 3A**). Thus platinum-resistant models may be divided into ones that respond to paclitaxel monotherapy and ones that are paclitaxel resistant as well as platinum-resistant.

Ascites-derived PDXO models from platinum-resistant patients (PDXO 2 and PDXO 80), as well as solid tumor-derived PDXO 80 exhibited resistance to all treatment conditions, showing no significant reduction in viability in response to therapy (**Figure 3B, C**).

However PDXO2 derived from solid tumors was sensitivity to paclitaxel and the combination treatment (**Figure 3C**). This demonstrates the inherent tumor heterogeneity in these samples.

In general, the consistent responses observed between ascites-derived PDOs and PDXOs reinforce their validity as reliable models for screening novel therapies and predicting patient treatment outcomes.

### Platinum-Refractory Models Show Resistance to Treatment

All three organoid models derived from the same platinum-refractory patient, PDO 1 (**Figure 4A**), PDXO 1 derived from ascites (**Figure 4B**), and PDXO 1 from solid tumors (**Figure 4C**), showed no response to carboplatin, paclitaxel, or the combination of carboplatin and paclitaxel, consistent with the treatment-refractory nature of this patient (**Figure 4A**).

Overall, our results suggest that ovarian cancer organoid models, either derived directly from the patient or first established as PDX in mice, are an equally good indicator of patient clinical response to standard chemotherapy. Furthermore, differences in response to single-agent paclitaxel might subdivide platinum-resistant patients potentially predicting their responsiveness to second-line paclitaxel therapy. This capability to forecast reactions to subsequent treatments could enhance therapeutic decision-making and optimize treatment sequences for resistant cancer cases [27, 42]. This study’s findings demonstrate that PDXO models faithfully reflect the varied responses of patients to chemotherapy, including significant reduction in viability for platinum-sensitive cases and carboplatin resistance in platinum-resistant or refractory cases. While PDOs provide valuable insights into patient-specific chemotherapy responses, the key advantage of using PDXOs is that they are a renewable source of patient-derived cancer cells, allowing for testing of novel treatments as they become available. By incorporating PDXOs into research, we can extend the utility of each patient sample, enabling a comprehensive investigation of potential therapies and facilitating the development of novel personalized treatment strategies.

## 4. DISCUSSION

Developing effective precision-targeted therapies necessitates preclinical models that represent the diverse morphological and molecular profiles of ovarian cancer tumors. Traditional models, particularly cancer cell lines grown as monolayers, fall short in replicating the genetic heterogeneity, morphology, and growth characteristics of human tumors, leading to limited predictive power for clinical outcomes [43]. Rapid growth of human cancer cells as a monolayer provides a strong selection for clonal outgrowth [44]. PDXs grown in immunocompromised mice better maintain the heterogeneity of the primary tumor and model 3D tumor growth, vascularization, and metastasis [13, 45, 46]. Given the time-consuming and costly nature of *in vivo* studies, *in vitro* drug screening with organoids is both beneficial and cost-effective, narrowing down candidate therapies to the most promising ones for subsequent *in vivo* testing. Our study highlights the potential of PDX-derived organoids as suitable preclinical models.

PDOs have been successfully cultured from many primary tumor types including breast cancer [47], lung [48], colon [49, 50], gastrointestinal [27], prostate [38, 51, 52], bladder [53, 54], ovarian [37, 55], endometrial [37, 56], and liver cancer [57]. These PDOs have shown promise in predicting patient responses to treatment. For example, the response of PDOs from patients with metastatic gastrointestinal cancer and colorectal cancer correlated with patient responses to anticancer agents observed in clinical trials [27, 58]. PDXOs have been successfully tested in other cancer types, including breast cancer where they provided insights into drug responses and therapy optimization [33-35] and in endometrial cancer, PDXOs were used to predict responses to FGFR inhibitors, showing potential in guiding treatment strategies [59]. However, their application in OC has been limited, making this study’s comparison of PDXOs and PDOs in OC particularly significant.

Uniquely, in our study, we developed organoids from both primary patient samples and PDX samples, from ascites fluid or solid tumors, to compare the drug response between these models and correlate the *in vitro* responses to the patient’s clinical response. Our findings demonstrate that both PDOs and PDXOs derived from platinum-sensitive OC patients exhibited a significant reduction in cell viability, of around 50%, when treated with carboplatin, paclitaxel, or their combination. This consistent response across different types of organoids mirrors previous patient-derive organoid studies where a reduction in cell viability correlates with a clinical platinum-sensitive response [29, 60, 61]. The ability of PDXOs to replicate these responses highlights their utility for preclinical drug testing and personalized medicine.

All organoid models from platinum-resistant and refractory cases were resistant to carboplatin monotherapy and the combination therapy, but varied in response to single agent paclitaxel. Future studies will use these PDXOs to test a broad range of second line therapies in order to tailor more effective treatment strategies for those not responding to first-line platinum-based therapies [61]. PDOs and PDXOs showed a similarity in response to treatments, indicating that sensitivity to standard chemotherapy of the primary cancer cells was maintained during establishment of the PDX, in agreement with previous PDX studies [12, 62] and during subsequent development of the PDXOs. Here we established paired PDOs and PDXOs (derived from ascites and solid tumors) from five patient samples, two from platinum-sensitive cases, two from platinum-resistant cases, and one from a platinum-refractory case. Our comparison of organoids derived from the same patient suggests that certain PDXOs may be more or less sensitive to individual chemotherapies compared to the primary PDO. For instance, although all of the organoid cultures derived from platinum-sensitive patient 20 were sensitive to single agent carboplatin and the combination standard treatment, both PDXO20 models were less sensitive to single agent carboplatin compared to the PDO (average PDXO viability -30% versus -50% in the PDO), suggesting a clonal bias in this PDX towards resistance. Platinum-resistant patient 2 exhibited the most differential responses to treatment among the different cultures. PDO2 was resistant to carboplatin and combination therapy, but sensitive to paclitaxel monotherapy (average -17% viability). PDXO2 derived from ascites was resistant to all treatments, while PDXO2 derived from a solid tumor, was more sensitive to single agent paclitaxel (average -27% viability) and combination therapy (average -16% viability). This suggests that different organoid cultures are sampling the heterogeneity of the patient cancer cells. None of the patient 2 organoid cultures showed organoid viability with single agent carboplatin of less than -30% indicating that the patient 2 cancer cells are primarily platinum-resistant. Overall, PDXOs retained their platinum-sensitivity or resistance supporting the utility of these models for preclinical testing. The variability in sensitivity among organoid models does indicate that testing multiple organoid models derived from different tumors will better represent patient cancer cell heterogeneity.

Here we directly compared PDOs derived from primary patient cancer cells and PDXOs derived from PDXs to demonstrate that these two patient-derived models show similar responses to standard chemotherapy. PDOs require less starting material and so can be established for a larger number of patients and are ready to assay for drug response in less than a month. However, while several independent laboratories have successfully established PDO living biobanks for OC [58, 63], a limited supply of primary tumor samples can be quickly depleted, and PDOs often exhibit reduced viability when thawed. PDX tumor tissue can be easily cryopreserved and serves as a renewable source for PDXOs, and that PDXs can be passaged into new NSG mice. Thus, PDXOs from the same PDX can be used in subsequent drug response studies, allowing comparisons of drug response across many studies. Importantly, PDXOs can be used as an initial *in vitro* screen to predict treatment response of PDX models in order to choose appropriate PDX with diverse responses to test *in vivo* [38] or to pre-screen larger drug libraries to identify candidate treatments to optimize personalized therapies *in vivo*.

While PDOs and PDXOs are a considerable improvement over cancer cell lines, PDOs still do not fully replicate the tumor microenvironment (TME). In our recent studies, we have been able to co-culture PDXOs with macrophages to examine the effect of macrophages on therapeutic response [64]. Enhancing organoid cultures with additional TME components, such as immune cells and stromal elements, could improve their representational accuracy [65]. Additionally, while organoids are valuable for short-term drug screening, they cannot capture long-term treatment responses, such as the development of drug resistance. This is why paired *in vitro* PDXO and *in vivo* PDX studies can be so powerful. Future studies will aim to integrate promising therapies identified in organoid models into *in vivo* testing using our humanized PDX (huPDX) models [13] to correlate with organoid results.

## 5. CONCLUSIONS

In conclusion, this study shows that response to chemotherapy in both PDO and PDXO models closely correlates with patient clinical responses. By comparing these models, we highlight PDXOs as a valuable, renewable platform for long-term drug screening and therapeutic testing. PDXOs capture the heterogeneity of OC, particularly when multiple organoid models can be derived from each patient, and extends the use of patient-derived samples, making them ideal for personalized medicine. Their reliable performance across treatments reinforces their potential in developing precision-targeted therapies, offering a versatile tool for advancing oncology research.

## List of abbreviations

(OC): Ovarian cancer
(PDOs: patient-derived organoids
(PDXOs: patient-derived xenograft organoids
(huPDX: humanized patient-derived xenografts
(PTX: paclitaxel
(CBDCA: carboplatin

## ACKNOWLEDGMENT

Funding was provided by a UNMCCC RAC Grant (Steinkamp), UNM Pathology Department and UNMCCC as startup funds (Dr. Steinkamp) and a UNMCCC Graduate Student Support Grant for P. Nikeghbal. We gratefully acknowledge use of the UNMCCC Animal Models (Irina Lagutina, PhD and Lillian Fitzpatrick), Fluorescence Microscopy (Diane S. Lidke, PhD and Michael Paffett, PhD), and the TSI Correlative Science Lab Shared Resource (Sarah Adams, MD and Chelsea Gregory) as well as the NIH P30CA118100 grant that supports the UNMCCC and these shared resources. Kim Leslie, PhD (UNM), and Kristina Thiel, PhD, (The University of Iowa) were instrumental in helping our lab with protocols to develop PDOs. As always, Dr. Diane S. Lidke provided helpful scientific advice. Additional technical assistance was provided by Shayna Lucero, Rachel Grattan, and Aubrey C. Gibson.

## AUTHOR CONTRIBUTIONS

Conceptualization: Parisa Nikeghbal and Mara P. Steinkamp; methodology: Parisa Nikeghbal and Mara Steinkamp; software: statistical analysis was performed using GraphPad Prism by Parisa Nikeghbal; validation: Parisa Nikeghbal and Mara P. Steinkamp; formal analysis: Parisa Nikeghbal; investigation: Parisa Nikeghbal, Dorsa Zamanian; resources: Mara P. Steinkamp; data curation: Parisa Nikeghbal, Dorsa Zamanian, Danielle Burke; writing—original draft preparation: Parisa Nikeghbal; review and editing: Mara P. Steinkamp, Danielle Burke; visualization: Parisa Nikeghbal; supervision: Mara P. Steinkamp; project administration: Mara P. Steinkamp; funding acquisition: Mara Steinkamp. All authors have read and agreed to the published version of the manuscript.

## INFORMED CONSENT STATEMENT

Malignant ascites fluid and fresh primary OC tumor pieces were collected from consenting OC patients undergoing cytoreductive surgery at the University of New Mexico Comprehensive Cancer Center under a UNM HSC IRB-approved protocol (INST 1509), as detailed in Steinkamp et al, 2023.

## REFERENCES

1. Webb, P.M. and S.J. Jordan, Global epidemiology of epithelial ovarian cancer. Nature Reviews Clinical Oncology, 2024. 21(5): p. 389–400.

2. Armstrong, D.K., et al., Ovarian cancer, version 2.2020, NCCN clinical practice guidelines in oncology. Journal of the National Comprehensive Cancer Network, 2021. 19(2): p. 191–226.

3. Kemp, Z. and J. Ledermann, Update on first-line treatment of advanced ovarian carcinoma. Int J Womens Health, 2013. 5: p. 45–51.

4. Lheureux, S., M. Braunstein, and A.M. Oza, Epithelial ovarian cancer: evolution of management in the era of precision medicine. CA: a cancer journal for clinicians, 2019. 69(4): p. 280–304.

5. Matulonis, U.A., et al., Ovarian cancer (Primer). Nature Reviews: Disease Primers, 2016. 2(1).

6. Davis, A., A.V. Tinker, and M. Friedlander, “Platinum resistant” ovarian cancer: what is it, who to treat and how to measure benefit? Gynecologic oncology, 2014. 133(3): p. 624–631.

7. Barakat, R.R., M. Markman, and M. Randall, Principles and practice of gynecologic oncology. 2009: Lippincott Williams & Wilkins.

8. Gorski, J.W., et al., Utilizing patient-derived epithelial ovarian cancer tumor organoids to predict carboplatin resistance. Biomedicines, 2021. 9(8): p. 1021.

9. Newtson, A., et al., Abstract B032: Use of patient-derived organoids to model tumor evolution in response to chemotherapy. Clinical Cancer Research, 2024. 30(5_Supplement): p. B032–B032.

10. Scott, C.L., et al., Patient-derived xenograft models to improve targeted therapy in epithelial ovarian cancer treatment. Frontiers in oncology, 2013. 3: p. 295.

11. Colombo, P.-E., et al., Ovarian carcinoma patient derived xenografts reproduce their tumor of origin and preserve an oligoclonal structure. Oncotarget, 2015. 6(29): p. 28327.

12. Liu, J.F., et al., Establishment of patient-derived tumor xenograft models of epithelial ovarian cancer for preclinical evaluation of novel therapeutics. Clinical Cancer Research, 2017. 23(5): p. 1263–1273.

13. Steinkamp, M.P., et al., Humanized patient-derived xenograft models of disseminated ovarian cancer recapitulate key aspects of the tumor immune environment within the peritoneal cavity. Cancer Research Communications, 2023. 3(2): p. 309–324.

14. Pizzi, M. and G. Inghirami, Patient-derived tumor xenografts of lymphoproliferative disorders: are they surrogates for the human disease? Current Opinion in Hematology, 2017. 24(4): p. 384–392.

15. Moro, M., et al., Establishment of patient derived xenografts as functional testing of lung cancer aggressiveness. Scientific Reports, 2017. 7(1): p. 6689.

16. Senkowski, W., et al., A platform for efficient establishment and drug-response profiling of high-grade serous ovarian cancer organoids. Developmental Cell, 2023. 58(12): p. 1106–1121. e7.

17. Kapalczynska, M., et al., 2D and 3D cell cultures–a comparison of different types of cancer cell cultures. Archives of medical science, 2018. 14(4): p. 910–919.

18. Świerczewska, M., et al., The response and resistance to drugs in ovarian cancer cell lines in 2D monolayers and 3D spheroids. Biomedicine & Pharmacotherapy, 2023. 165: p. 115152.

19. Patra, B., et al., Carboplatin sensitivity in epithelial ovarian cancer cell lines: The impact of model systems. PLoS One, 2020. 15(12): p. e0244549.

20. Ciucci, A., et al., Preclinical models of epithelial ovarian cancer: practical considerations and challenges for a meaningful application. Cell Mol Life Sci, 2022. 79(7): p. 364.

21. Yee, C., et al., Three-dimensional modelling of ovarian cancer: From cell lines to organoids for discovery and personalized medicine. Frontiers in Bioengineering and Biotechnology, 2022. 10: p. 836984.

22. Maru, Y., et al., Efficient use of patient-derived organoids as a preclinical model for gynecologic tumors. Gynecologic oncology, 2019. 154(1): p. 189–198.

23. Tiriac, H., et al., Organoid profiling identifies common responders to chemotherapy in pancreatic cancer. Cancer discovery, 2018. 8(9): p. 1112–1129.

24. Yan, H.H., et al., A comprehensive human gastric cancer organoid biobank captures tumor subtype heterogeneity and enables therapeutic screening. Cell stem cell, 2018. 23(6): p. 882–897. e11.

25. de Witte, C.J., et al., Patient-derived ovarian cancer organoids mimic clinical response and exhibit heterogeneous inter-and intrapatient drug responses. Cell reports, 2020. 31(11).

26. Ooft, S.N., et al., Patient-derived organoids can predict response to chemotherapy in metastatic colorectal cancer patients. Science translational medicine, 2019. 11(513): p. eaay2574.

27. Vlachogiannis, G., et al., Patient-derived organoids model treatment response of metastatic gastrointestinal cancers. Science, 2018. 359(6378): p. 920–926.

28. Jabs, J., et al., Screening drug effects in patient-derived cancer cells links organoid responses to genome alterations. Molecular systems biology, 2017. 13(11): p. 955.

29. Kopper, O., et al., An organoid platform for ovarian cancer captures intra-and interpatient heterogeneity. Nature medicine, 2019. 25(5): p. 838–849.

30. Swan, H.A., et al., personalized medicine: a CLIA-certified high-throughput drug screening platform for ovarian cancer. Cancer Research, 2018. 78(13_Supplement): p. 1619–1619.

31. Larsen, B.M., et al., A pan-cancer organoid platform for precision medicine. Cell reports, 2021. 36(4).

32. Nanki, Y., et al., Patient-derived ovarian cancer organoids capture the genomic profiles of primary tumours applicable for drug sensitivity and resistance testing. Scientific reports, 2020. 10(1): p. 12581.

33. Scherer, S.D., et al., Breast cancer PDxO cultures for drug discovery and functional precision oncology. STAR protocols, 2023. 4(3): p. 102402.

34. Guillen, K.P., et al., A human breast cancer-derived xenograft and organoid platform for drug discovery and precision oncology. Nature cancer, 2022. 3(2): p. 232–250.

35. Navarro-Yepes, J., et al., Abemaciclib is effective in palbociclib-resistant hormone receptor–positive metastatic breast cancers. Cancer research, 2023. 83(19): p. 3264–3283.

36. Thorel, L., et al., Comparative analysis of response to treatments and molecular features of tumor-derived organoids versus cell lines and PDX derived from the same ovarian clear cell carcinoma. Journal of experimental & clinical cancer research, 2023. 42(1): p. 260.

37. Bi, J., et al., Successful patient-derived organoid culture of gynecologic cancers for disease modeling and drug sensitivity testing. Cancers, 2021. 13(12): p. 2901.

38. Colling, K.E., et al., Multiplexed live-cell imaging for drug responses in patient-derived organoid models of cancer. JoVE (Journal of Visualized Experiments), 2024(203): p. e66072.

39. Rose, P.G., Ovarian cancer recurrence: Is the definition of platinum sensitivity modified by PARPi, bevacizumab or other intervening treatments?: A clinical perspective. Cancer Drug Resistance, 2022. 5(2): p. 415.

40. Markman, M. and W. Hoskins, Responses to salvage chemotherapy in ovarian cancer: a critical need for precise definitions of the treated population. Journal of Clinical Oncology, 1992. 10(4): p. 513–514.

41. St Laurent, J. and J.F. Liu, Treatment Approaches for Platinum-Resistant Ovarian Cancer. Journal of Clinical Oncology, 2024. 42(2): p. 127–133.

42. Gao, J., et al., Promising preclinical patient-derived organoid (PDO) and xenograft (PDX) models in upper gastrointestinal cancers: progress and challenges. BMC cancer, 2023. 23(1): p. 1205.

43. Weeber, F., et al., Tumor organoids as a pre-clinical cancer model for drug discovery. Cell chemical biology, 2017. 24(9): p. 1092–1100.

44. Porter, S.N., et al., Lentiviral and targeted cellular barcoding reveals ongoing clonal dynamics of cell lines in vitro and in vivo. Genome Biology, 2014. 15(5): p. R75.

45. Lawrence, M.G., et al., The future of patient-derived xenografts in prostate cancer research. Nature Reviews Urology, 2023. 20(6): p. 371–384.

46. Hidalgo, M., et al., Patient-derived xenograft models: an emerging platform for translational cancer research. Cancer discovery, 2014. 4(9): p. 998–1013.

47. Sachs, N., et al., A living biobank of breast cancer organoids captures disease heterogeneity. Cell, 2018. 172(1): p. 373–386. e10.

48. Sachs, N., et al., Long-term expanding human airway organoids for disease modeling. The EMBO journal, 2019. 38(4): p. e100300.

49. Jung, P., et al., Isolation and in vitro expansion of human colonic stem cells. Nature medicine, 2011. 17(10): p. 1225–1227.

50. Sato, T., et al., Long-term expansion of epithelial organoids from human colon, adenoma, adenocarcinoma, and Barrett’s epithelium. Gastroenterology, 2011. 141(5): p. 1762–1772.

51. Chua, C.W., et al., Single luminal epithelial progenitors can generate prostate organoids in culture. Nature cell biology, 2014. 16(10): p. 951–961.

52. Gil, V., et al., HER3 is an actionable target in advanced prostate cancer. Cancer research, 2021. 81(24): p. 6207–6218.

53. Lee, S.H., et al., Tumor evolution and drug response in patient-derived organoid models of bladder cancer. Cell, 2018. 173(2): p. 515–528. e17.

54. Cai, E.Y., et al., A bladder cancer patient-derived xenograft displays aggressive growth dynamics in vivo and in organoid culture. Scientific reports, 2021. 11(1): p. 4609.

55. Gonzalez Bosquet, J., et al., Prediction of chemo-response in serous ovarian cancer. Molecular cancer, 2016. 15: p. 1–15.

56. Bi, J., et al., Advantages of tyrosine kinase anti-angiogenic cediranib over bevacizumab: cell cycle abrogation and synergy with chemotherapy. Pharmaceuticals, 2021. 14(7): p. 682.

57. Broutier, L., et al., Human primary liver cancer–derived organoid cultures for disease modeling and drug screening. Nature medicine, 2017. 23(12): p. 1424–1435.

58. Van de Wetering, M., et al., Prospective derivation of a living organoid biobank of colorectal cancer patients. Cell, 2015. 161(4): p. 933–945.

59. Sengal, A.T., et al., Endometrial cancer PDX-derived organoids (PDXOs) and PDXs with FGFR2c isoform expression are sensitive to FGFR inhibition. npj Precision Oncology, 2023. 7(1): p. 127.

60. Tao, M., et al., Developing patient-derived organoids to predict PARP inhibitor response and explore resistance overcoming strategies in ovarian cancer. Pharmacological research, 2022. 179: p. 106232.

61. Xiang, D., et al., Building consensus on the application of organoid-based drug sensitivity testing in cancer precision medicine and drug development. Theranostics, 2024. 14(8): p. 3300.

62. Liu, Y., et al., Patient-derived xenograft models in cancer therapy: technologies and applications. Signal Transduction and Targeted Therapy, 2023. 8(1): p. 160.

63. Wang, E., et al., Patient-derived organoids (PDOs) and PDO-derived xenografts (PDOXs): new opportunities in establishing faithful pre-clinical cancer models. Journal of the National Cancer Center, 2022. 2(4): p. 263–276.

64. Nikeghbal, P., et al., The Influence of Macrophages within the Tumor Microenvironment on Ovarian Cancer Growth and Response to Therapies. bioRxiv, 2025: p. 2025.01.29.635532.

65. Neal, J.T., et al., Organoid modeling of the tumor immune microenvironment. Cell, 2018. 175(7): p. 1972–1988. e16.

