## Supplementary material for "Organoid Models Established from Primary Tumors and Patient-Derived Xenograft Tumors Reflect Platinum Sensitivity of Ovarian Cancer Patients": Figures

**Supplemental Figure 1. Growth of PDOs derived from primary malignant ascites of high grade OC primary patients**

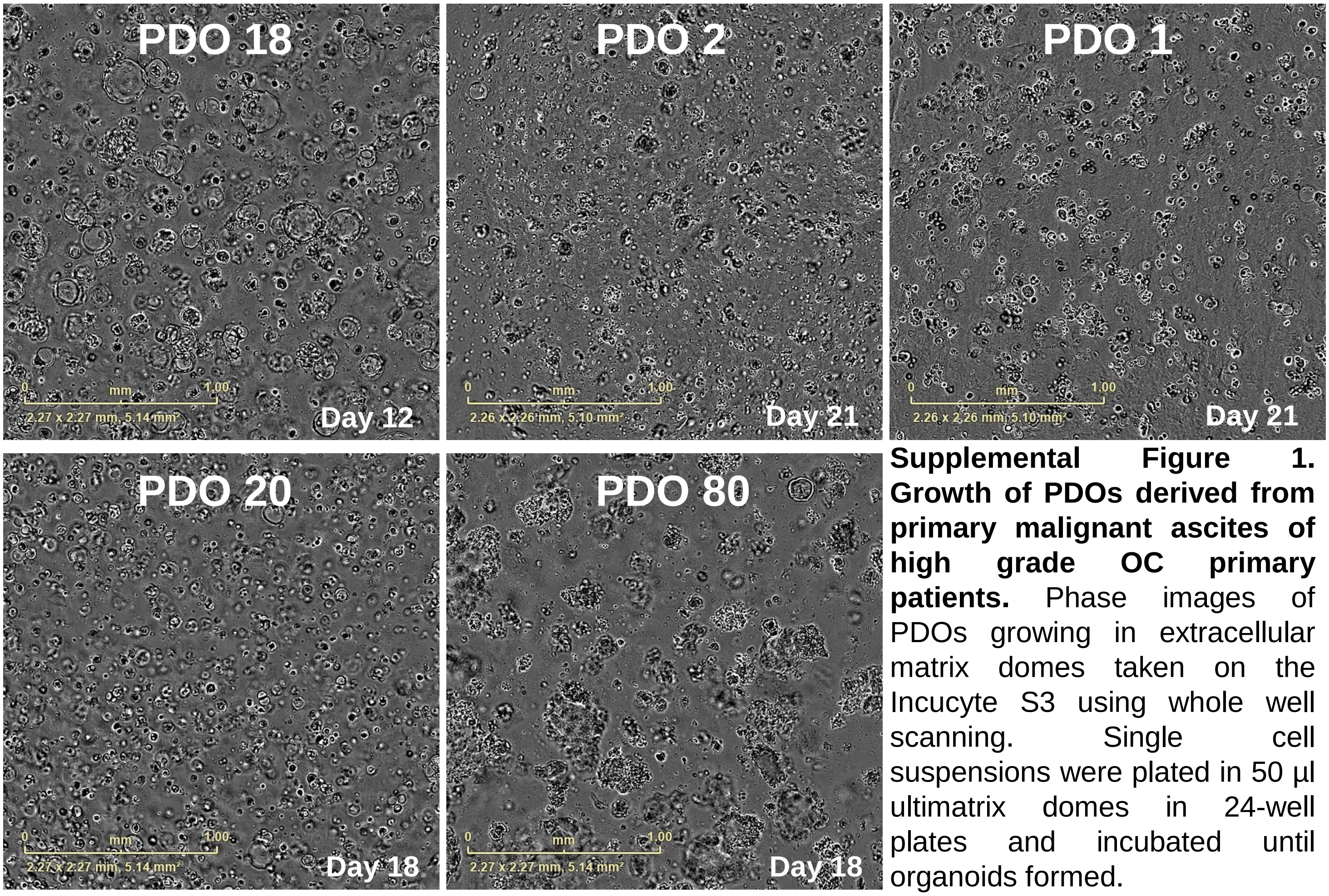


**Supplemental Figure 1. Growth of PDOs derived from primary malignant ascites of high grade OC primary patients.** Phase images of PDOs growing in extracellular matrix domes taken on the Incucyte S3 using whole well scanning. Single cell suspensions were plated in 50 µl ultimatrix domes in 24-well plates and incubated until organoids formed.

**Supplemental Figure 2. Growth of PDXOs derived from PDX ascites samples shows similarities to PDOs formed from primary ascites.**

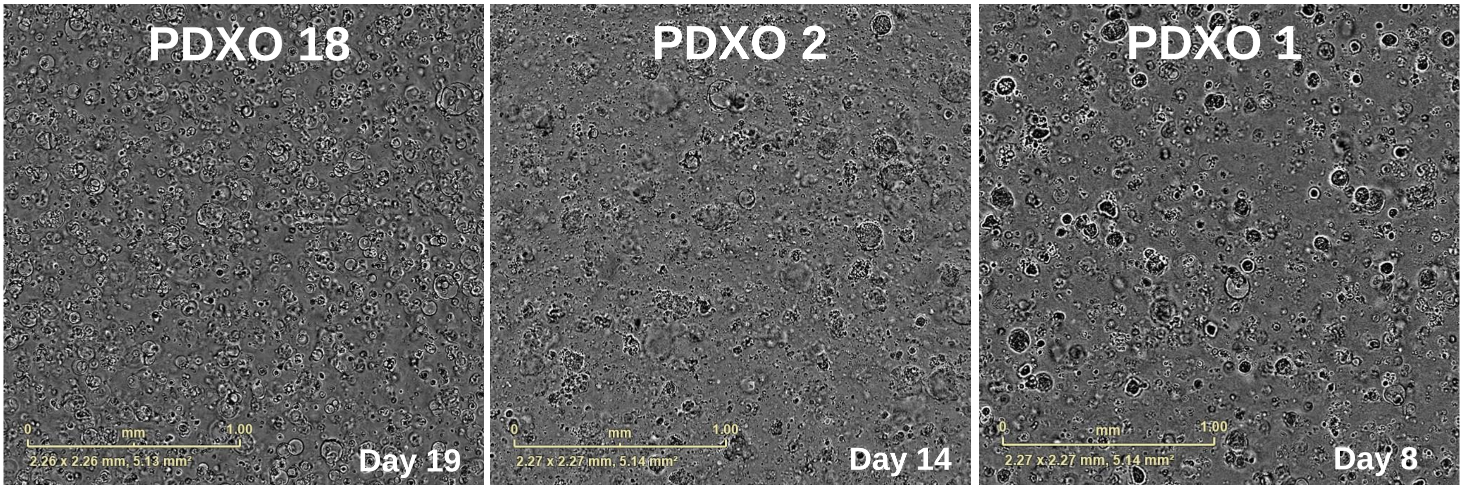


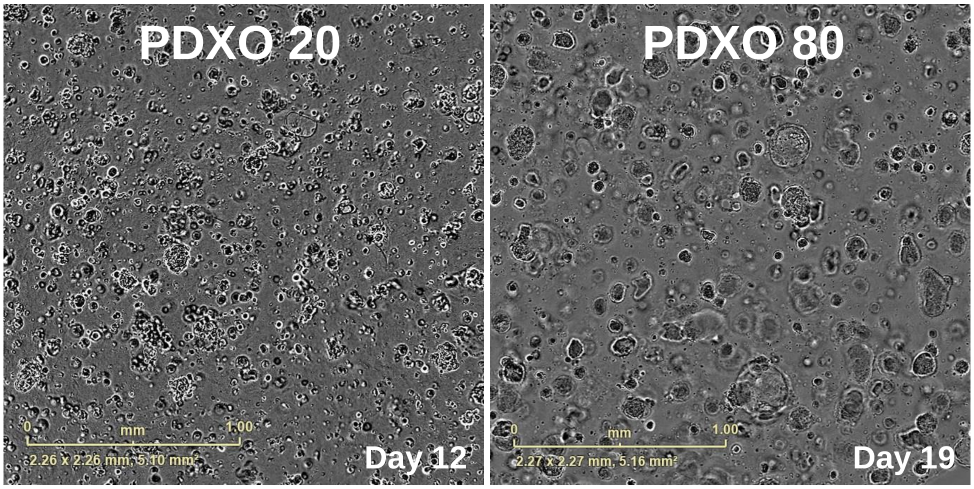


**Supplemental Figure 2. Growth of PDXOs derived from PDX ascites samples shows similarities to PDOs formed from primary ascites.** Phase images of four ascites-derived PDXOs growing in ultimatrix domes. Single cell suspensions were plated in 50 µl ultimatrix domes in 24-well plates and incubated until organoids formed.

**Supplemental Figure 3. PDXOs derived from PDX solid tumors**

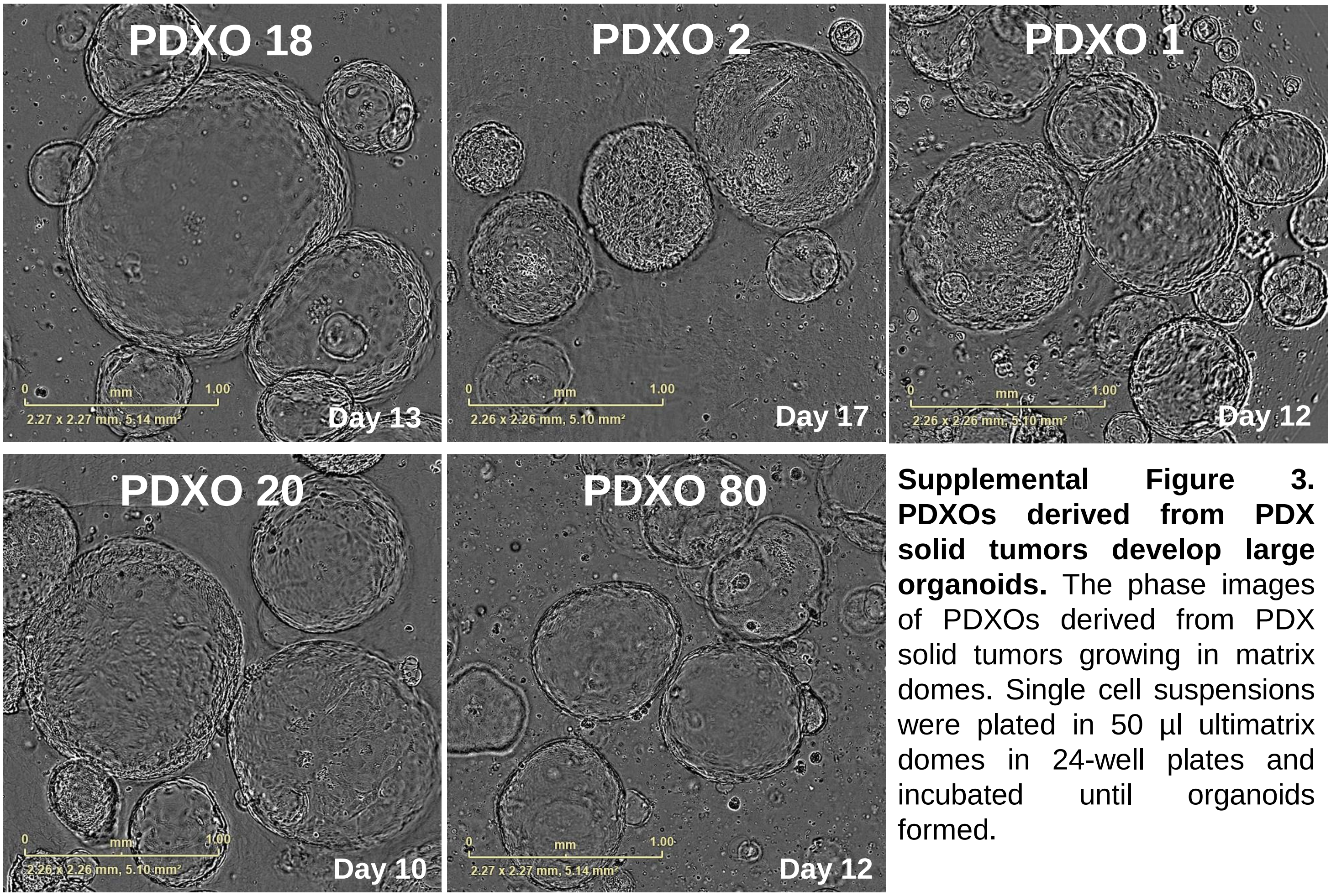


**Supplemental Figure 3. PDXOs derived from PDX solid tumors develop large organoids.** The phase images of PDXOs derived from PDX solid tumors growing in matrix domes. Single cell suspensions were plated in 50 µl ultimatrix domes in 24-well plates and incubated until organoids formed.

**Supplemental Figure 4. Raw Viability Data Comparing Untreated and 10 µM Afatinib Treated Organoid Models.**

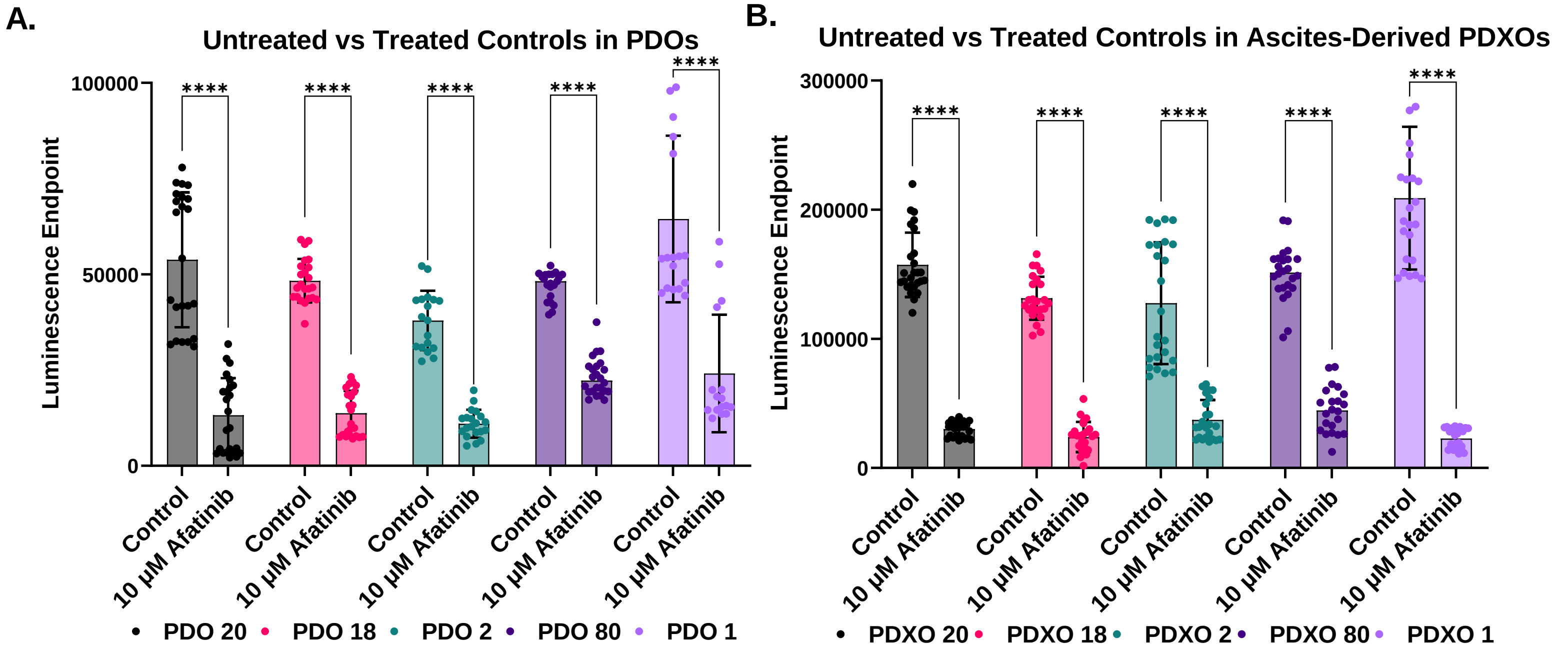


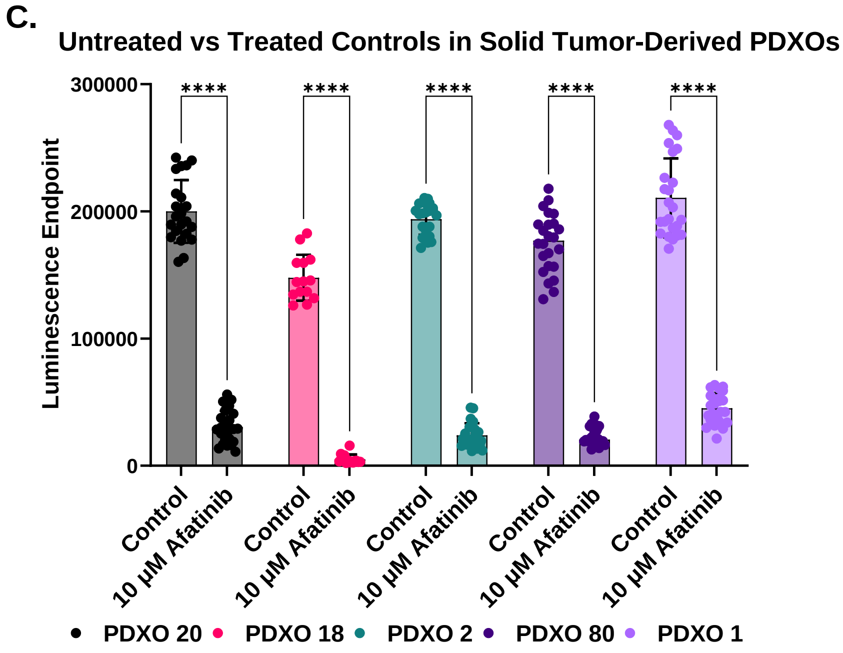


**Supplemental Figure 4. Raw Viability Data Comparing Untreated and 10 µM Afatinib Treated Organoid Models.** Viability measurements for untreated and 10 µM Afatinib-treated samples are shown for each model. **A.** PDOs. **B**. PDXOs derived from ascites, and **C.** PDXOs derived from solid tumor. Untreated organoids have significantly higher luminescence compared to the 10 µM afatinib control indicating that all untreated control organoids were metabolically active. based on two biological replicates with 6 or more technical replicates per experiment. Statistical significance is marked by asterisks (**** $p<0.0001$), assessed via one-way ANOVA.
